# Seasonal and depth variations in diet composition and dietary overlap between three native killifish of an emblematic tropical-mountain lake: Lake Titicaca (Bolivia)

**DOI:** 10.1101/635821

**Authors:** Erick Loayza

**Affiliations:** Unidad de Limnología, Instituto de Ecología, Universidad Mayor de San Andrés, La Paz, Bolivia

## Abstract

Lake Titicaca (∼3800 m a.s.l.), an emblematic tropical-mountain ecosystem is the major source of fish for people on the Altiplano. The Andean killifish genus *Orestias*, represent an important resource for local fisheries in Lake Titicaca. It has been suggested that exist an effect of segregation in the Lake Titicaca in order to avoid competition for food resource between native fish species, due most of *Orestias* species share the littoral habitat, which is now also share with introduced species. Such scenario increases the pressure for food resource. Here I examined the gut content of *O. luteus, O. agassizii* and *O. mulleri* (Cuvier & Valenciennes, 1846) from a bay of Lake Titicaca during rainy (April) and dry season (July) with the predominance method, frequency of occurrence and numerical percentage to describe the diet and dietary overlap between these native fish. I also applied a PERMANOVA test in order to determine diet variations related to depth and seasonally, as well as the Levins and Pianka’s index to test diet breadth and dietary overlap respectively. 396 gut contents were evaluated, identifying a high frequency of amphipods and molluscs in the three *Orestias* native species. Diet breadth revelled a selectivity for a few preys and the composition of the diets was influenced mainly by depth, followed by seasonality (PERMANOVA, P = <0.05). Dietary overlapping between *O. luteus* and *O. agassizii* was evidenced in the rainy season. During the dry season, the three species undergone dietary overlapping. This study provided a detail knowledge on the diet variations of native species in Lake Titicaca, especially for *Orestias mulleri*, a little-known species. Here I also discussed the importance of the amphipods as a food resource in Lake Titicaca not only for fish community, but for the food web in general. The seasonal and depth diet variations here discussed are relevant for fisheries management and conservation and could be used to guide aquaculture development in Lake Titicaca.

## 1 INTRODUCTION

The Altiplano is one of the largest high plateaus in the world containing the Lake Titicaca, the largest navigable water body in the world (3809 m a.s.l.), and also the most important water resource of the Andean region. Lake Titicaca represents the major source of fish for ∼3 million people on the Altiplano, between native and introduced fish. Waters of Lake Titicaca are mainly oligotrophic, with almost constant light and temperature conditions and permanently hyperhaline due to the geographical characteristics and the lack of strong seasonality on the region (Dejoux & Iltis, 1992). Nevertheless, it is not clear if this lack of seasonality has an influence on the behaviour or foraging strategies of the native ichthyofaunal, represented mainly for *Orestias* (Valenciennes, 1839) one of the endemic genus of the Altiplano (Dejoux & Iltis, 1992; Vila, Pardo & Scott, 2007).

*Orestias* have 23 species described for Lake Titicaca, although only a few are recognized (Dejoux & Iltis, 1992; Vila *et al.*, 2007; Ibañez *et al.*, 2014). It has been suggested that exist an effect of segregation in the habitat, reason why exist such morphological variability in this genus (Lauzanne, 1982; Loubens, 1989; Dejoux & Iltis, 1992; Maldonado *et al.*, 2009). *Orestias* are an important piece in the trophic network in Lake Titicaca, however, their diet descriptions are based mainly on general observations and not on specific studies (Ibañez *et al.*, 2014). In addition, most of *Orestias* species have benthic habits and share the littoral habitat with juveniles of pejerrey (*Odontesthes bonariensis*, Valenciennes, 1835) an introduced species (Monroy *et al.*, 2014).

*Orestias agassizii* (Cuvier & Valenciennes, 1846) and *Orestias luteus* (Cuvier & Valenciennes, 1846) are the *Orestias* with most economically relevant for local fisheries. They coexist throughout the lake and are frequently found in the littoral zone near to the shore. However, *O. agassizii* is capable of being in littoral and pelagic zones, while *O. luteus* inhabits benthic zone, where it coexists with *Orestias mulleri* (Cuvier & Valenciennes, 1846), which is considered a bentopelagic fish (Monroy *et al.*, 2014). Nowadays, there is a lack of knowledge about the trophic interactions, diet breadth and other aspects of feeding ecology of *Orestias*, due the studies on these fish were focused on morphological and taxonomic analysis (Ibañez *et al.*, 2014; Guerrero-Jiménez *et al.*, 2017).

*Orestias* usually inhabit littoral zone in Lake Titicaca, as well as smaller sizes of introduced species such as trout (*Oncorhynchus mykiss*, Walbum, 1792) and pejerrey (*Odontesthes bonariensis*, Valenciennes, 1835) so they belong to the same trophic level (Monroy *et al.*, 2014). Therefore, there is a niche overlap and competition for food resource are very likely, however there are no studies that prove this hypothesis. It is well known that feeding is a non-linear behaviour with many scaled gradients, such as time (i.e., time of year), space (i.e., change through depth), morphology (i.e., morphology of the prey or size of the predator) or other biological attributes (Saikia, 2016).

Future environmental changes are inevitable, especially in relation to new environmental problems such as climate change and the pressures of invasive species, which represent a common threat to the native fish populations. This can affect the functions of an ecosystem and trophic relationships (predator-prey interactions), which are a very important component of studies at the ecosystem level, particularly because species can modify their diet in response to these changes. Therefore, here I describe the diet, their breadth and dietary overlap of three native species (*O. agassizii, O. luteus* and *O. mulleri*) that coexist in a bay of Lake Titicaca, a tropical-mountain ecosystem. Further, I evaluate the diet variations in relation to depth and seasonality.

## 2 METHODS

### 2.1 Study area

Lake Titicaca is the largest freshwater lake in South America, with 8559 km^2^ area and located at 3810 m.a.s.l. It is divided into two sub-basins: Lago Mayor, which reaches 285 m maximum depth, and Lago Menor, with a maximum 40 m depth (Dejoux & Iltis, 1992). Although there is a lack of seasonality on the region, exist a marked increase in rainfall (between December and March) and a dry season (between May and August) (Myers *et al.*, 2000; Vila *et al.*, 2007).

This study focused on Toke Pucuro Bay, near the small town of Achacachi (Figure 1), which, like most of the shores on Lago Mayor, has three types of habitats: 1) the pelagic zone (i.e., open waters of the lake) with abundant cladocerans and other zooplankton; 2) the benthic zone (i.e., near bottom area) rich in molluscs and amphipods and 3) the coastal zone characterized by being rich in macrophytes such as totoras *(Schoenoplectus californicus* ssp. t*atora*), juncus (*Juncus articus* ssp. *andicola*), and other genera such as *Chara, Potamogeton, Myriophyllum, Nitella* and *Ruppia* in which a large number of amphipods are found (Dejoux & Iltis, 1992; Lauzanne, 1992; Vila *et al.*, 2007). These vegetation area represents an important area of feeding and reproduction for fish in the lake (Lauzanne, 1992).

**Figure 1.**
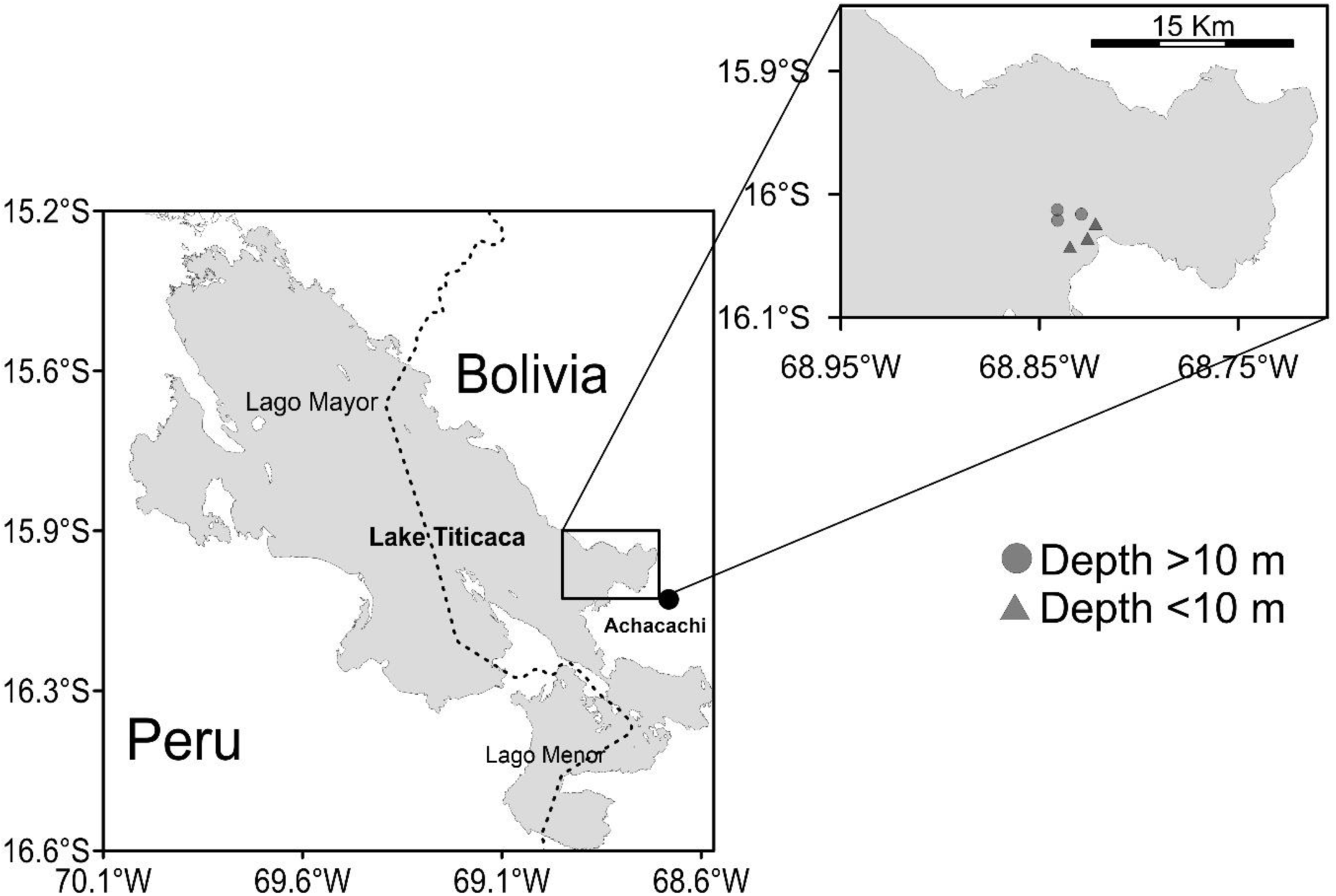
Sampling location of shallow waters fishing sites (< 10 m) (grey triangle) and deep water (> 10 m) (grey circle) in Lake Titicaca in April and July 2018.

### 2.2 Fish sampling

Experimental gillnets of 12 panels of 11 mm to 110 mm openings were used, as well as gillnets (48 mm opening) from a local fisherman. Fish sampling were made at the end of the rainy season (April) and during the dry season (July) 2018. 3 shallow habitats with depth <10 m (9.1 m max. depth) and 3 pelagic habitats with depth > 10 m (21.4 m max. depth) were sampled evenly distributed in the study area. At the same time, 3 samples of benthic invertebrate collected at each fish sampling site were taken with an Eckman dredger to determine the composition of the possible fish prey. The samples of benthic invertebrate were fixed in 10% formalin and were identified at the highest taxonomic level possible.

Fish were identified at the species level, measured, weighed and euthanized in 96% ethanol (Metcalfe & Craig, 2011). Guts were removed *in situ* and fixed in 75% ethanol to avoid degradation of the gut contents. Fish total length (TL) were measured to the nearest 0.01 mm using a digital meter. Fish and gut were weighed to the nearest 0.01 g using a digital scale. Gut contents were examined with a microscope (X40) in which the identifiable parts of the organisms were considered as individuals and identified with the lowest taxon possible.

### 2.3 Analysis

Representativeness of the samplings was estimated using an accumulation curve randomized with respect to the number of gut contents reviewed. Prey diversity consumed by three *Orestias* native species studied was established by the Simpson index (Magurran, 2013):

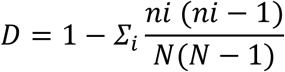

Where “*N*” is the total number of prey and “*ni*” is the number of individuals of prey “*i*” (Hurlbert, 1971). Prey richness (number of prey in the gut content) was calculated and the diet breadth using the standardized Levins index (Levins, 1974; Krebs, 1999):

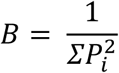

Where “*pi*” is the proportion of individuals of the “*i*” prey found in one of three study species. Levins index standardized (BA = B-1 / n-1) was applied to express the diet breadth in a scale that fluctuates between 0 and 1. Lower values than 0.60 are considered as a specialized diet using a low resource number, and values above 0.60 as a generalist (Krebs, 1999). Pianka’s symmetric index (1974) was measured to estimate the niche overlap in the diet composition between each species, depth and season (Guerrero *et al.*, 2015). It is considered a biologically significant overlap when the value of this index exceeds 0.6 (Pianka, 1974).

Diet composition was quantified by a semi-quantitative visual estimate of the prey abundance (zooplankton, amphipods, insects, macrophytes, algae, molluscs, ostracods, sediments, fish eggs and others), according to five categories: absent (0%), very rare (25%), rare (50%), abundant (75%) and very abundant (100%) following the modifications of the predominance method (Frost & Went, 1940; Tresierra Aguilar & Culquichicón Malpica, 1993). Frequency of occurrence (%FO) and numeric percentage (%N) (Hyslop, 1980; Zavala-Camin, 1996) of each species according to depth and season was expressed as:

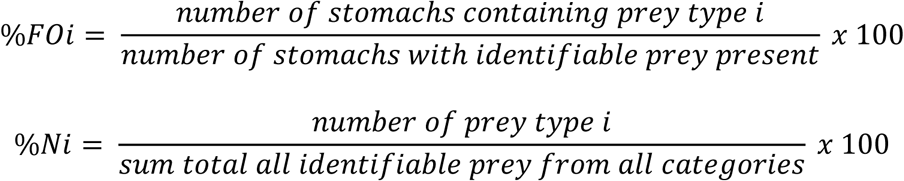

Gravimetric or volumetric measurements were not made, since the presence of sediment and detritus in gut content makes them unfeasible, as fractionation and different digestibility of each component diet could bias this measure (Cardona, 1991), as well as generating problems in the interpretation (Baker, Buckland & Sheaves, 2014; Buckland *et al.*, 2017).

An analysis of similarities (ANOSIM, α = 0.05) of the distances of Bray-Curtis with the abundances of benthic invertebrate with 9999 permutations was performed to test differences in the composition of the benthic preys between depths and season. To test intraspecific between depths and seasonal differences in diet composition, permutational multivariate analysis of variance (PERMANOVA) of the abundance of the gut content was applied, using the similarity of Bray-Curtis with 9999 permutations. Processing and analysis were performed in RStudio, version 1.1.453 (RStudio 2016) with R, version 3.4.0 (R Core Team, 2018) and the packages vegan (Oksanen *et al.*, 2018), spaa (Jinlong, 2016) and BiodiversityR (Kindt & Coe, 2005).

## 3 RESULTS

### 3.1 Benthic invertebrate composition in habitat

In total were recorded nineteen taxa of benthic invertebrate (Table 1). Taxa richness between depths was the same at the end of rainy season (April), slightly different during dry season (July). On the other hand, the abundance did not change between depths, but they were different between seasons. *Hyalella* spp. (40.32% at a depth < 10 m, 46.29% at a depth > 10 m) and Hydrobiidae (25.43% at a depth < 10 m, 16.20% at a depth > 10 m) abundances were higher during rainy season, whereas during dry season were *Hyalella* spp. (73.76% at a depth < 10 m, 64.19% at a depth > 10 m) and Hirudinea (10.40% at a depth < 10 m, 13.07% at a depth < 10 m).

**TABLE 1.**
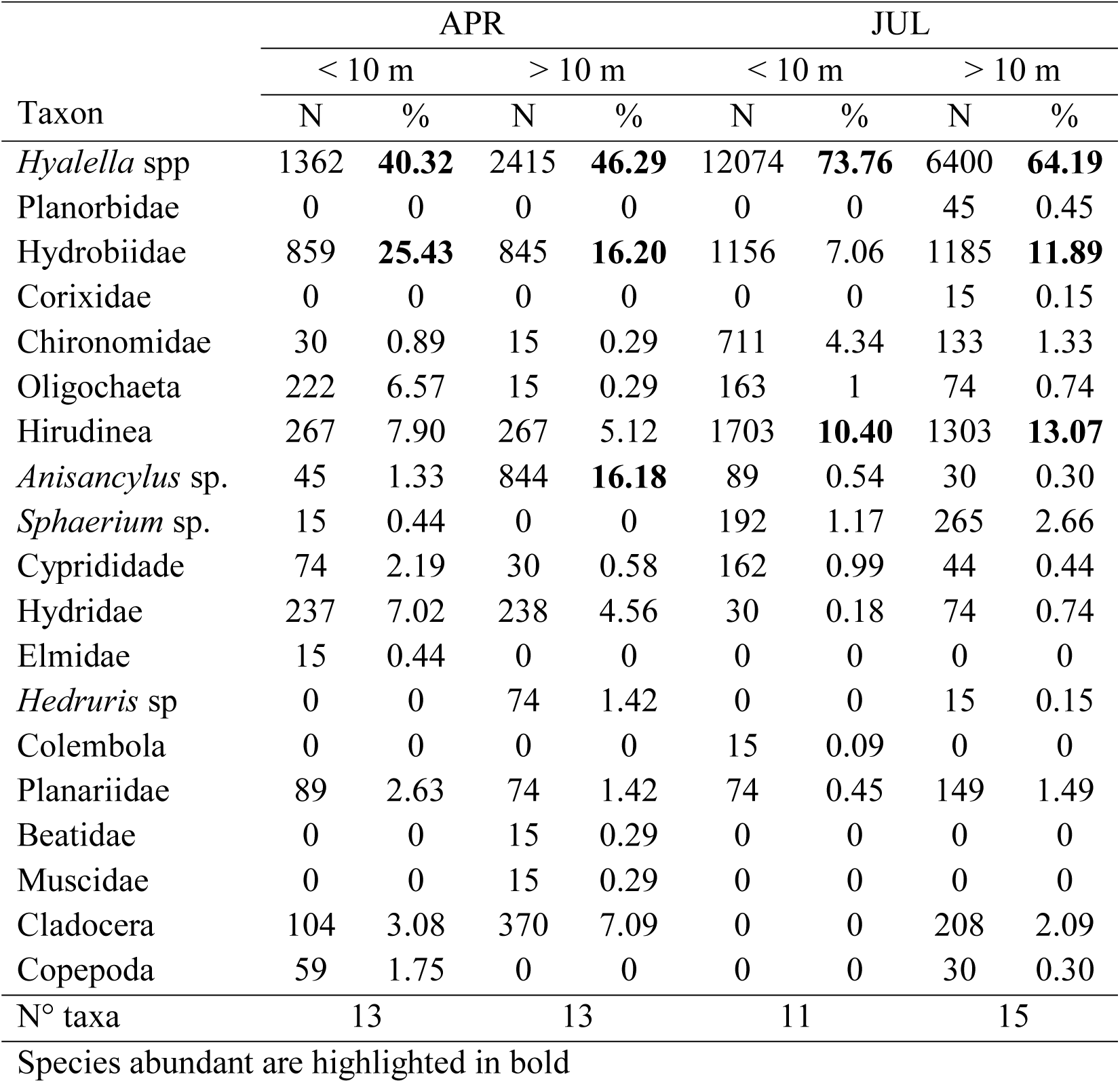
Abundance of benthic invertebrates in two different depths and seasons. End of the rainy season (APR), dry season (JUL) in Toke Pucuro Bay, Lake Titicaca.

| Taxon | APR |  |  |  | JUL |  |  |  |
| --- | --- | --- | --- | --- | --- | --- | --- | --- |
|  | < 10 m |  | > 10 m |  | < 10 m |  | > 10 m |  |
|  | N | % | N | % | N | % | N | % |
| *Hyalella* spp | 1362 | **40.32** | 2415 | **46.29** | 12074 | **73.76** | 6400 | **64.19** |
| Planorbidae | 0 | 0 | 0 | 0 | 0 | 0 | 45 | 0.45 |
| Hydrobiidae | 859 | **25.43** | 845 | **16.20** | 1156 | 7.06 | 1185 | **11.89** |
| Corixidae | 0 | 0 | 0 | 0 | 0 | 0 | 15 | 0.15 |
| Chironomidae | 30 | 0.89 | 15 | 0.29 | 711 | 4.34 | 133 | 1.33 |
| Oligochaeta | 222 | 6.57 | 15 | 0.29 | 163 | 1 | 74 | 0.74 |
| Hirudinea | 267 | 7.90 | 267 | 5.12 | 1703 | **10.40** | 1303 | **13.07** |
| *Anisancylus* sp. | 45 | 1.33 | 844 | **16.18** | 89 | 0.54 | 30 | 0.30 |
| *Sphaerium* sp. | 15 | 0.44 | 0 | 0 | 192 | 1.17 | 265 | 2.66 |
| Cyprididae | 74 | 2.19 | 30 | 0.58 | 162 | 0.99 | 44 | 0.44 |
| Hydriidae | 237 | 7.02 | 238 | 4.56 | 30 | 0.18 | 74 | 0.74 |
| Elmidae | 15 | 0.44 | 0 | 0 | 0 | 0 | 0 | 0 |
| *Hedruris* sp | 0 | 0 | 74 | 1.42 | 0 | 0 | 15 | 0.15 |
| Colembola | 0 | 0 | 0 | 0 | 15 | 0.09 | 0 | 0 |
| Planariidae | 89 | 2.63 | 74 | 1.42 | 74 | 0.45 | 149 | 1.49 |
| Beatidae | 0 | 0 | 15 | 0.29 | 0 | 0 | 0 | 0 |
| Muscidae | 0 | 0 | 15 | 0.29 | 0 | 0 | 0 | 0 |
| Cladocera | 104 | 3.08 | 370 | 7.09 | 0 | 0 | 208 | 2.09 |
| Copepoda | 59 | 1.75 | 0 | 0 | 0 | 0 | 30 | 0.30 |
| N° taxa | 13 |  | 13 |  | 11 |  | 15 |  |

Species abundant are highlighted in bold

Benthic invertebrate composition showed a significant difference between seasons (p <0.01) and the R value for depths comparison was close to 0, which indicates that benthic invertebrate composition was similar to each other (Table 2).

**TABLE 2.** Analysis of similarity (ANOSIM) of two pathways of the composition of aquatic invertebrates in different seasons and depths from the Bray-Curtis distances with the abundances of benthic invertebrates with 9999 permutations.

| Factor | R | *p* (perm) |
| --- | --- | --- |
| Season | 0.51215 | **0.0051** |
| Depth | -0.09028 | 0.7101 |

Significant P-values are highlighted in bold

### 3.2 Diet composition of three *Orestias* species

In total 396 gut contents were evaluated. 36 were empty (*Orestias luteus* = 21, *Orestias agassizii* = 13, *Orestias mulleri* = 2) and thus not analysed. Accumulation curve showed that the number of gut contents evaluated was adequate to make the inferences (Figure 2). To facilitate the analysis, the taxa Oligocheta, *Hydrozetes* sp. and Hirudinea were grouped in one category, named as “*Other*”, due to their low representation in gut contents.

**Figure 2.**
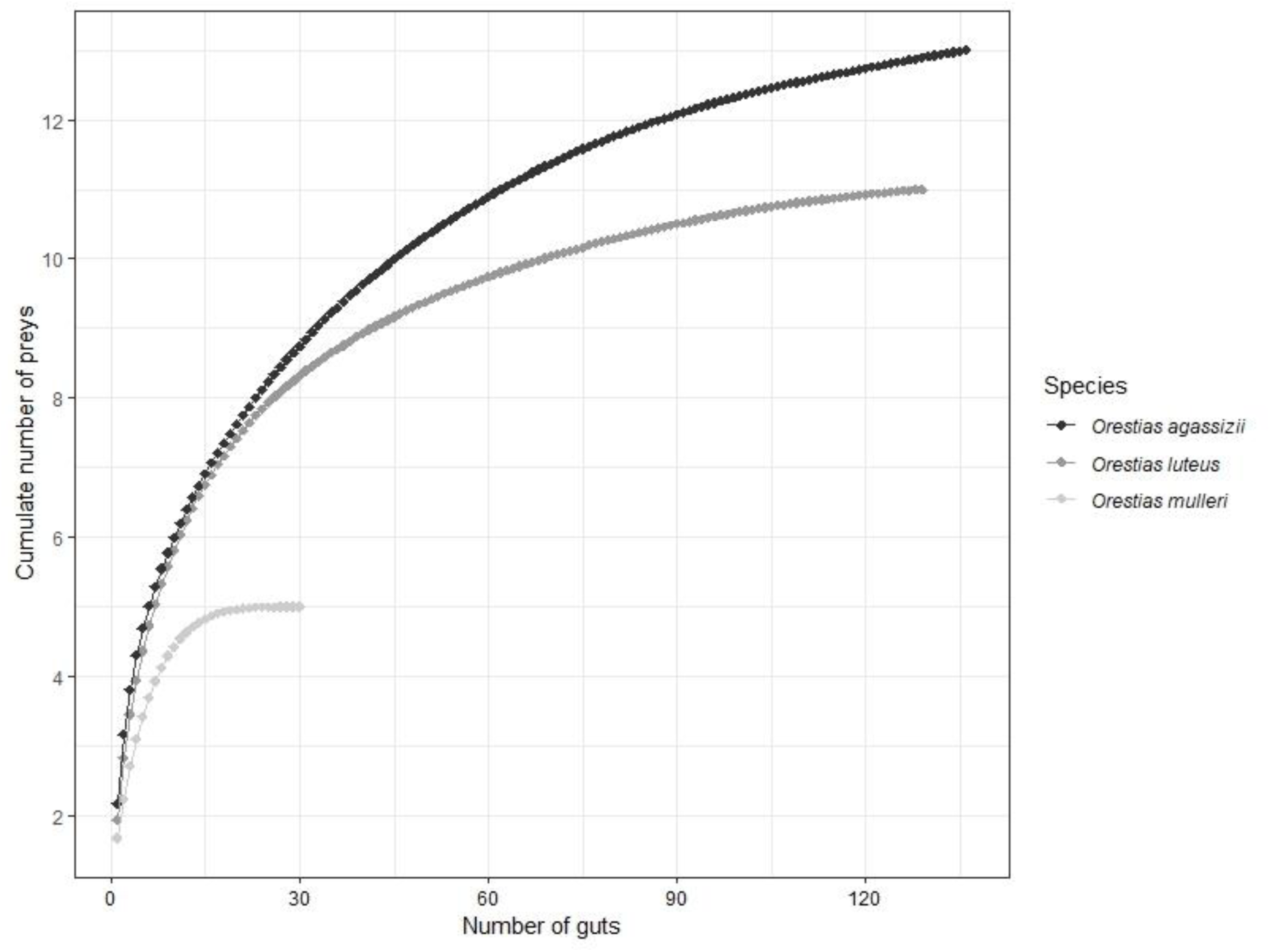
Preys accumulation curve with respect to the number of *Orestias* gut contents sampled in Lake Titicaca.

In general, the diet of these three *Orestias* species was based on amphipods, being the group most consumed (Figure 3). During rainy season, *O. luteus* showed intra-specific differences in their diet. The main prey in shallow waters were the amphipods (71.5%) and molluscs (62.7%) in waters with depths < 10 m. In contrast, during dry season its diet was based on amphipods (60.3% at a depth < 10 m and 76.8% at a depth > 10 m) and molluscs (28.5% at a depth < 10 m and 6.2% at a depth > 10 m). Prey diversity consumed (Simpson index) by this species was higher during rainy season, although diet breadth did not reflect such diversity (B_A_ <0.6) (Table 3).

**TABLE 3.** Diversity (D), prey richness (S), Levins index (B) and standardized Levins index (B_A_) of three *Orestias* species. End of the rainy season (APR), dry season (JUL).

| Species | Season | Depth | N | Simpson index (D) | S | Simpson index (D) per season | S per season | B season | B<sub>A</sub> season | Simpson index (D) per specie | S per specie | B Specie | B<sub>A</sub> Specie |
| --- | --- | --- | --- | --- | --- | --- | --- | --- | --- | --- | --- | --- | --- |
| *Orestias luteus* | APR | < 10 m | 63 | 0.44 | 10 | 0.58 | 10 | 2.40 | 0.16 | 0.53 | 11 | 2.06 | 0.11 |
|  |  | >10 m | 26 | 0.44 | 5 |  |  |  |  |  |  |  |  |
|  | JUL | < 10 m | 48 | 0.33 | 9 | 0.33 | 9 | 1.49 | 0.06 |  |  |  |  |
|  |  | >10 m | 20 | 0.34 | 8 |  |  |  |  |  |  |  |  |
| *Orestias agassizii* | APR | < 10 m | 47 | 0.58 | 11 | 0.71 | 12 | 3.47 | 0.22 | 0.69 | 13 | 2.93 | 16 |
|  |  | >10 m | 46 | 0.69 | 11 |  |  |  |  |  |  |  |  |
|  | JUL | < 10 m | 31 | 0.53 | 7 | 0.53 | 7 | 2.14 | 0.19 |  |  |  |  |
|  |  | >10 m | 60 | 0.53 | 7 |  |  |  |  |  |  |  |  |
| *Orestias mulleri* | APR | >10 m | 18 | 0.46 | 5 | 0.46 | 5 | 1.86 | 0.22 | 0.65 | 5 | 2.88 | 0.47 |
|  | JUL | >10 m | 14 | 0.52 | 3 | 0.52 | 3 | 2.07 | 0.54 |  |  |  |  |

**Figure 3.**
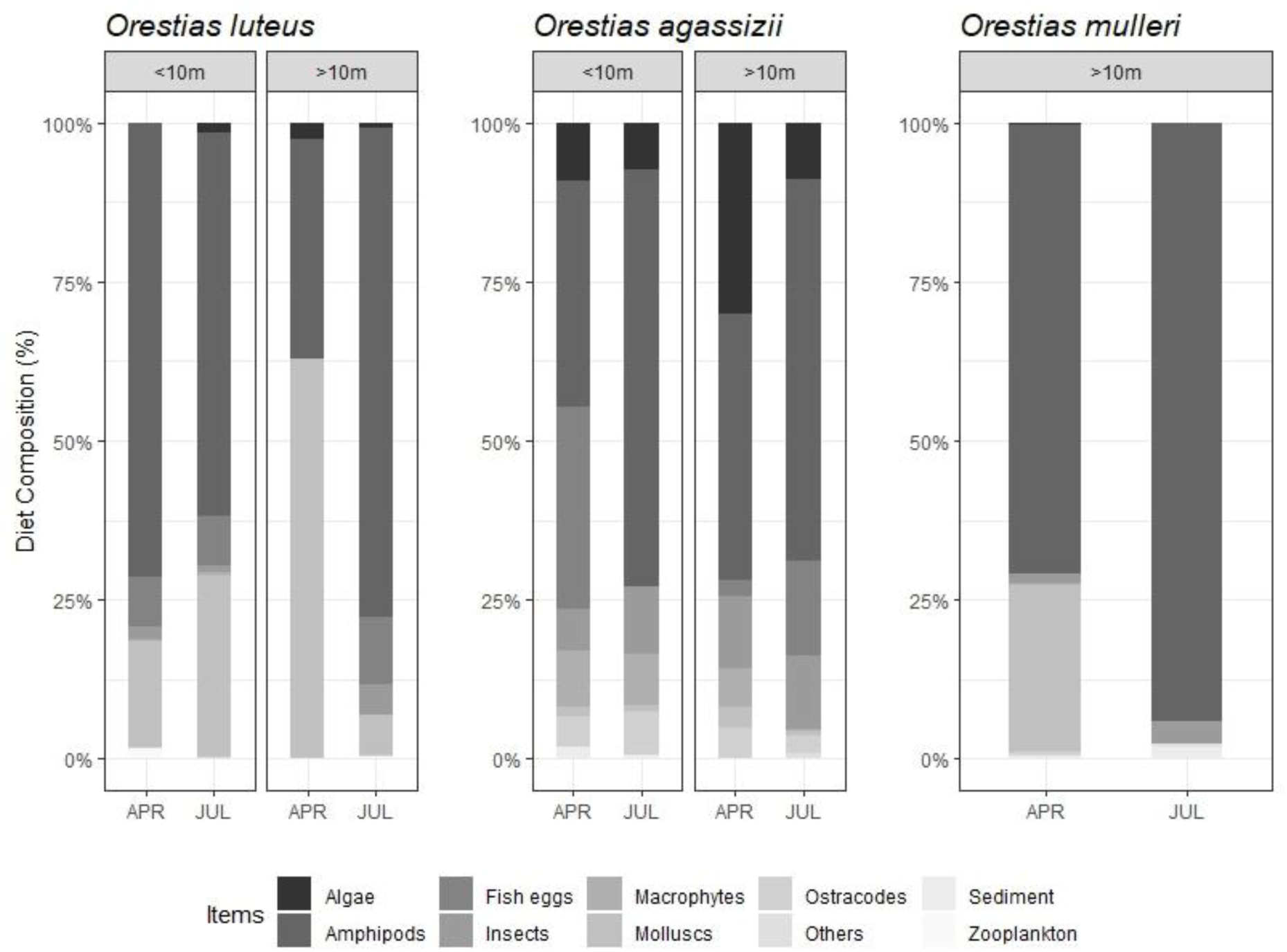
Predominance in diet composition of *Orestias luteus* (N = 157), *Orestias agassizii* (N = 184) and *Orestias mulleri* (N = 32) from Lake Titicaca during end of rainy season (APR) and dry season (JUL) in 2018.

*O. agassizii* has amphipods as its main prey, however, it was able to take advantage of a larger number of resources (D = 0.71, S = 12) (Table 3). During rainy season it was fed on fish eggs (31.7%) in shallow water, and algae (30.1%) in deep water. On the other hand, during dry season *O. agassizii* fed of amphipods (65.7%) and insects (10.3%) and in areas with depths < 10 m, it was also fed on fish eggs (14.9%). Diversity of prey consumed was lower during this season, with a reduced trophic spectrum (B_A_ = 0.19).

*O. mulleri*, a species that was only found in deep waters (< 10 m), amphipods (70.6%) and molluscs (26.2%) were the main food items during rainy season. In contrast, for dry season, there was an almost exclusive feeding of amphipods (94%). *O. mulleri* showed prey diversity indexes similar between both seasons (D = 0.46, D = 0.52 respectively), with a trophic breadth higher than the other *Orestias* species (0.47). Further, low prey richness (S = 5) in both seasons (S = 5, S = 3; respectively) was observed (Table 3).

### 3.3 Seasonal and depth variations in diet composition of three *Orestias* species

Due to its low representativeness (<3%) both frequency of occurrence analysis, and numerical percentage analysis, Oligocheta, *Hydrozetes* sp., Hirudinea, *Anisancylus* sp. and sediments were grouped into a single category named “*Occasional prey*”. *O. luteus,* during rainy season in shallow waters, fed on *Hyalella* spp. whose frequency of occurrence (%FO) comprised 48.4% and 74.9% in numerical percentage (%N). Hydrobiidae had a frequency of 14.3% and Cladocera with 14.9% N (Figure 4 and 5). At higher depths, their diet was based on Hydrobiidae with 48.4%FO and 67.6%N, followed by *Hyalella* spp. with 37.3%FO and 21.6%N (Figure 4 and 5). During dry season *Hyalella* spp. represented a frequency of occurrence 35.7% and 50% for each depth, and a high numerical percentage of 75.7% and 80.3%.

**Figure 4.**
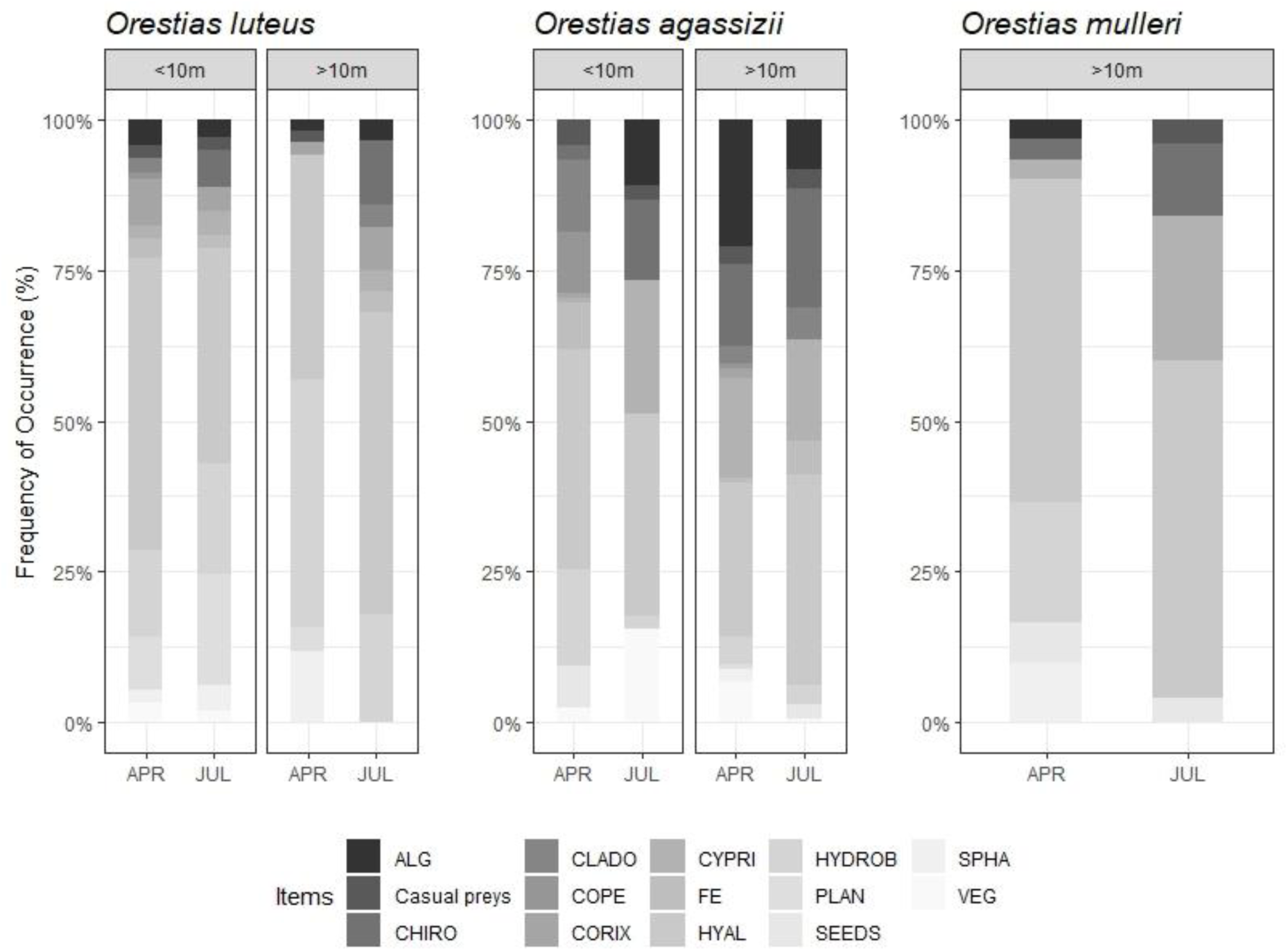
Frequency of occurrence (%FO) of prey taxa in diets of *Orestias luteus* (N = 157), *Orestias agassizii* (N = 184) and *Orestias mulleri* (N = 32) from Lake Titicaca during end of rainy season (APR) and dry season (JUL) in 2018. Preys: algae (ALG), Chironomidae (CHIRO), Cladocera (CLADO), Copepoda (COPE), Corixidae (CORIX), Cyprididae (CYPRI), fish eggs (FE), *Hyalella* spp. (HYAL), Hydrobiidae (HYDROB), Planorbidae (PLAN), macrophyte seeds (SEEDS), *Sphaerium* sp. (SPHA), vegetation (VEG).

**Figure 5.**
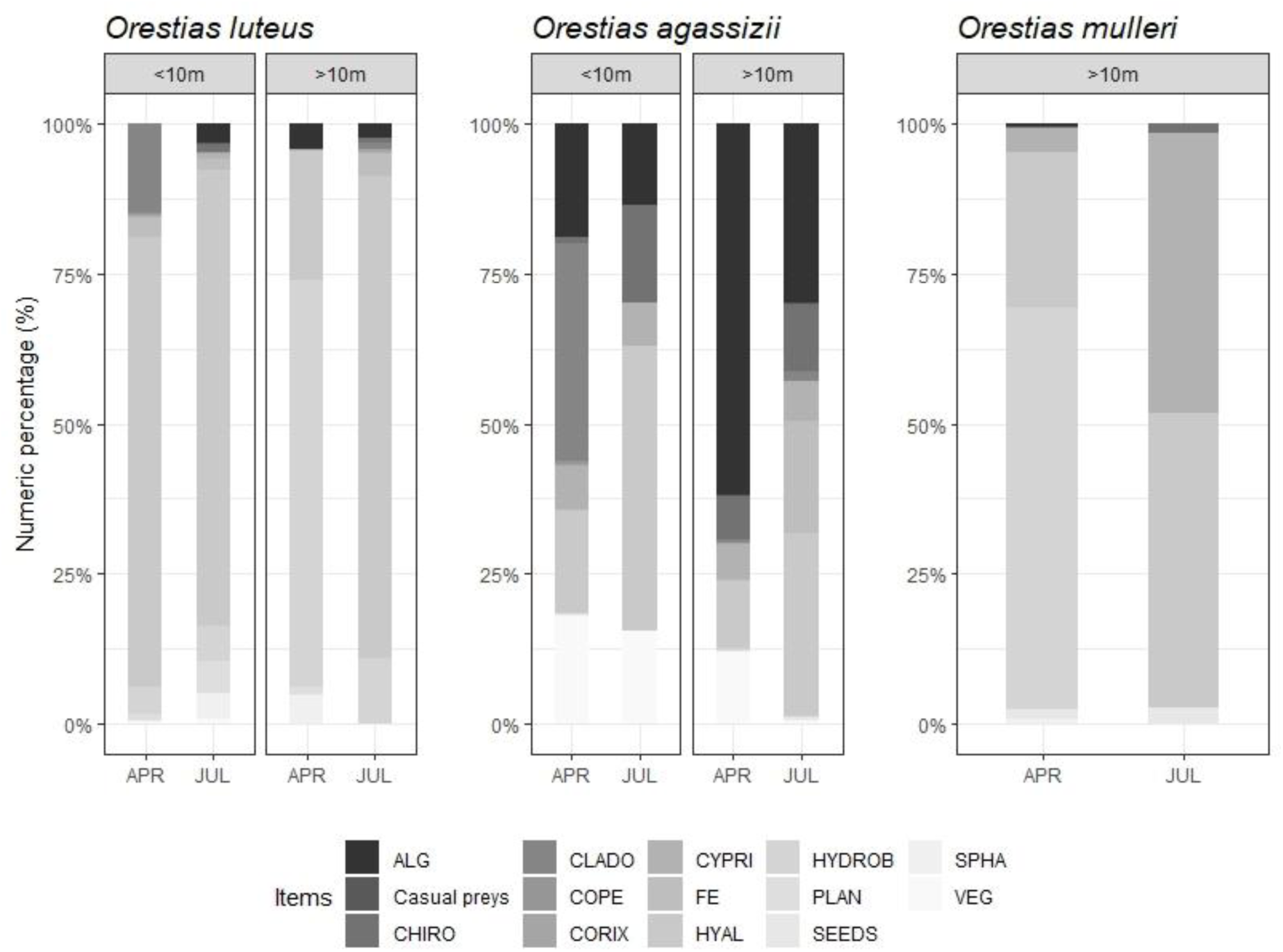
Numeric percentage (%N) of prey taxa in diets of *Orestias luteus* (N = 157), *Orestias agassizii* (N = 184) and *Orestias mulleri* (N = 32) from Lake Titicaca during end of rainy season (APR) and dry season (JUL) in 2018. Preys: algae (ALG), Chironomidae (CHIRO), Cladocera (CLADO), Copepoda (COPE), Corixidae (CORIX), Cyprididae (CYPRI), fish eggs (FE), *Hyalella* spp. (HYAL), Hydrobiidae (HYDROB), Planorbidae (PLAN), macrophyte seeds (SEEDS), *Sphaerium* sp. (SPHA), vegetation (VEG).

For *O. agassizii,* during rainy season at a depth < 10 m, *Hyalella* spp. had a frequency of occurrence of 36.4% and Hydrobiidae (16.1%) (Figure 4). Instead, at a depth > 10 m, amphipods represented 25.6%FO and algae 21.1%FO. On the other hand, the numerical percentage for *O. agassizii* was dominated by Cladocera in shallow waters (36.1%) and algae (61.8%) in deeper waters (Figure 5).

In *O. agassizii* gut content during dry season, the main consumed prey was *Hyalella* spp., which reached 33.3%FO and 35.1%FO, for each depth range. The importance of Cyprididae (22.2%FO) at a depth < 10 m, and Chironomidae (19.8%FO) at a depth > 10 m, increase during this season. Same patron is observed for amphipods in the numerical percentage, where they represent 47.2%N at a depth < 10 m of the *O. agassizii* gut content, and 30.3%N at a depth > 10 m. It is also remarkable that the intake of fish eggs (18.7%) increase.

*O. mulleri* was fed more frequently of *Hyalella* spp. during rainy season (53.3%FO) but with a higher numerical percentage of Hydrobiidae (67%N). Something similar was observed during dry season, where *Hyalella* spp. had a 56%FO and 48.9%N, followed by Cyprididae (46.7%N) (Figure 5).

### 3.4 Intraspecific variation in diet composition and dietary overlap of three *Orestias* species

Feeding habits of *O. luteus* and *O. agassizii* showed intra-specific variations in relation to depth, but are more influenced by the season (Table 4). PERMANOVA test showed a significant difference in feeding habits in relation to the interaction of the season with the depth for both species. In contrast, these habits were relatively consistent at both seasons for *O. mulleri*. Further, Pianka’s index indicate a total overlap between *O. luteus* and *O. agassizii* at the end of rainy season (Figure 6), which increases during dry season. Overlap was higher among all fish species and in both depths during dry season. *O. agassizii* and *O. luteus* in shallow waters had a higher overlap (0.94), there being a complete overlap between *O. agassizii* and *O. mulleri* at a depth > 10 m, followed by *O. luteus* with *O. mulleri* at the same depth range (Figure 7). Pianka’s index suggest that dietary overlap is higher between the three species at both depths during the dry season.

**TABLE 4.** Results of PERMANOVA between different seasons and depths on the diet of three *Orestias* species

| Source of variation | df | SS | MS | Pseudo-F | R<sup>2</sup> | p (perm) |
| --- | --- | --- | --- | --- | --- | --- |
| *Orestias luteus* |  |  |  |  |  |  |
| Season | 1 | 0.680 | 0.67959 | 2.2769 | 0.01668 | **0.0390** |
| Depth | 1 | 1.479 | 1.47940 | 4.9565 | 0.03632 | **0.0002** |
| Season*Depth | 1 | 1.268 | 1.26807 | 4.2485 | 0.03113 | **0.0010** |
| Residuals | 125 | 37.310 | 0.29848 |  | 0.91587 |  |
| *Orestias agassizii* |  |  |  |  |  |  |
| Season | 1 | 0.675 | 0.67515 | 2.3523 | 0.01664 | **0.0214** |
| Depth | 1 | 1.197 | 1.19677 | 4.1697 | 0.0295 | **0.0007** |
| Season*Depth | 1 | 0.813 | 0.81342 | 4.8341 | 0.02005 | **0.008** |
| Residuals | 132 | 37.886 | 0.28701 |  | 0.93381 |  |
| *Orestias mulleri* |  |  |  |  |  |  |
| Season | 1 | 0.4811 | 0.48109 | 1.9276 | 0.06441 | 0.1122 |
| Residuals | 28 | 6.9883 | 0.24958 |  | 0.93559 |  |

Significant P-values are highlighted in bold

**Figure 6.**
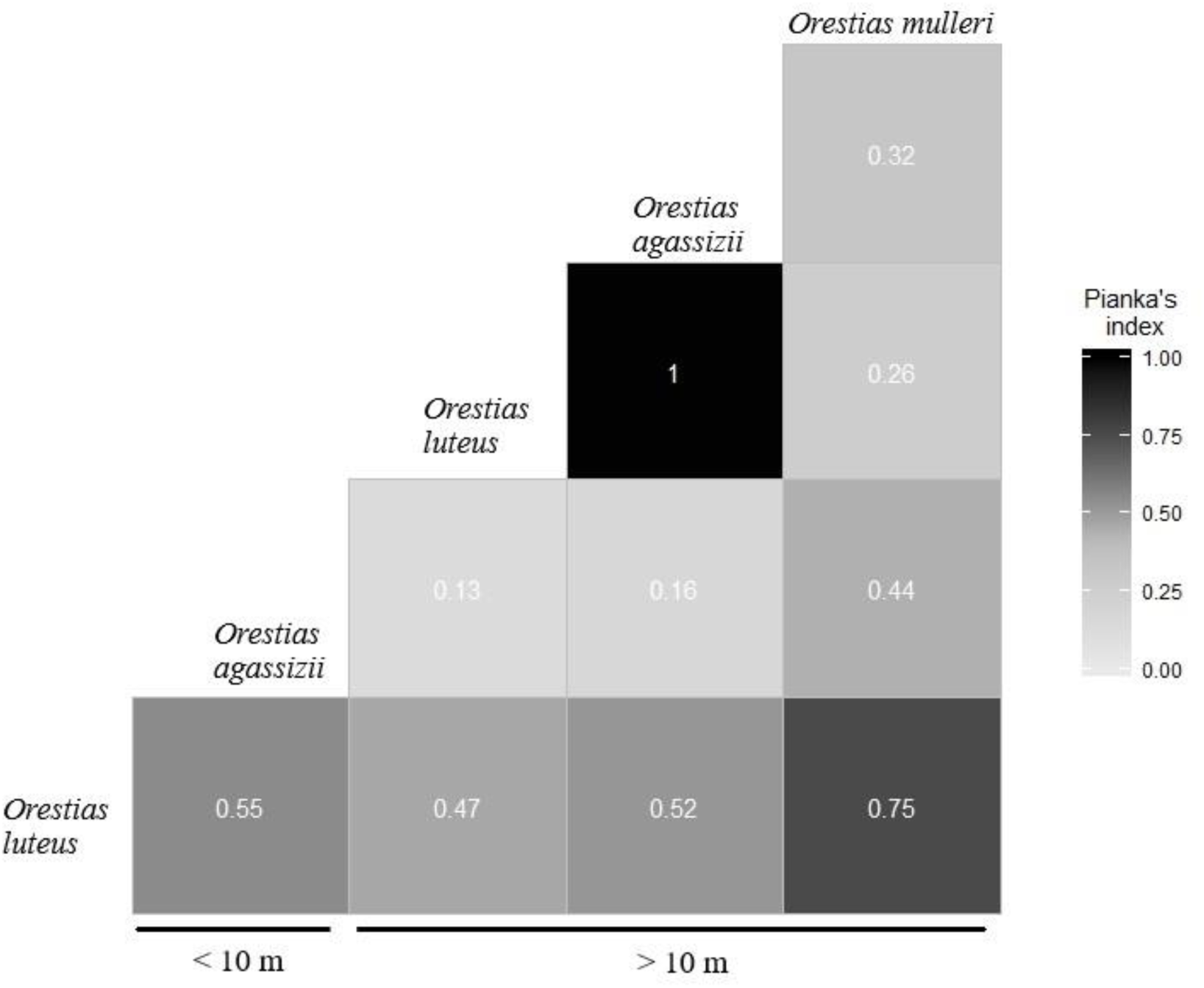
Dietary overlap between three *Orestias* species at the end of the rainy season (April) 2018.

**Figure 7.**
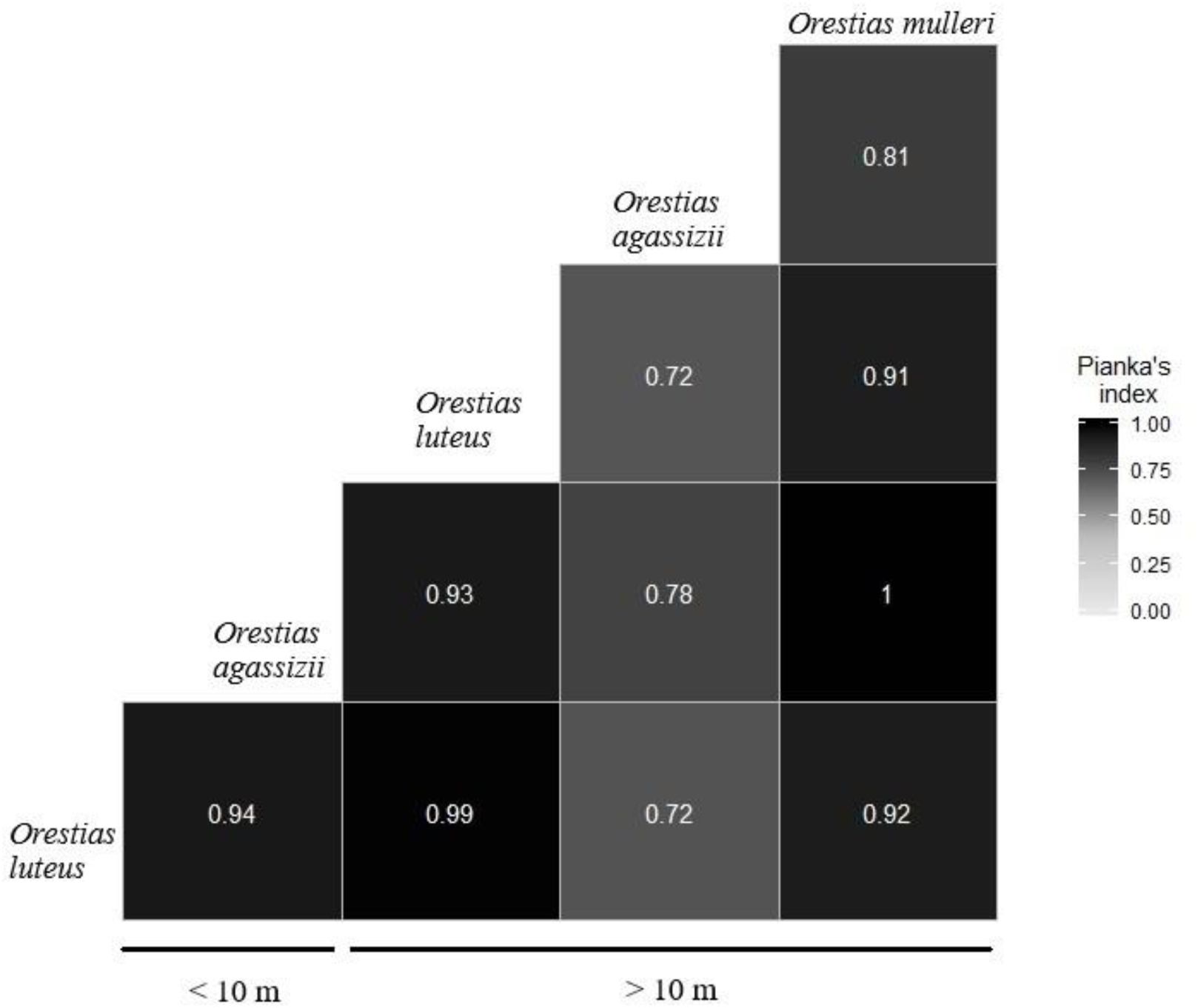
Dietary overlap between three *Orestias* species during dry season (July) 2018.

## 4 DISCUSSION

Benthic fauna in shallow bays at Lake Titicaca is mainly represented by molluscs and amphipods the former being the predominant group in the Characeas, whereas chironomids and amphipods dominate the macrophytes areas (Dejoux, 1992). These does not differ much in bare bottoms with deeps less than 20 m, where benthic fauna is higher in molluscs and amphipods (Dejoux, 1992). Both taxa represent an essential component in the biology of the lake due they perform an important role at trophic dynamics as well in energy transfers (Dejoux, 1992). During this study, the benthic fauna was largely composed of amphipods and molluscs (Hydrobiidae) and *Anisancylus* sp., which represent the main food resource for fish populations in the lake (Lauzanne, 1992; Vila *et al.*, 2007). The invertebrate composition did not change significantly with depth. However, during the dry season *Hyalella* spp. and Hirudinea had higher abundances than the rest of taxa, which was also observed in fish diets.

General observations indicated that *O. luteus* frequently inhabits shallow areas near to the shore of the lake, and usually feeds on aquatic insects and amphipods (Vila *et al.*, 2007). During this study it was observed that *O. luteus* feeds mainly of amphipods and molluscs (Hydrobiidae and Planorbidae), taking advantage of fish eggs as a resource in shallow waters. During dry season the patron remained the same, but the intake of fish eggs was higher, especially at a depth > 10 m. The intake of fish eggs by this species seems to be a frequent behaviour, also reported by Maldonado et al. (2009).

*O. agassizii* showed a varied diet, although similarly predominated by amphipods. *O. agassizii* is generally classified as a ubiquitous species, due to its ability to inhabit most lacustrine habitats (Lauzanne, 1992). Such ability was observed in the feeding habits, because during rainy season it was fed on zooplankton (Cladocera) at a depth < 10 m, and algae at a depth >10 m. In contrast, during dry season it was also fed on ostracods (Cyprididade), algae and vegetation (macrophytes) at a depth < 10 m, and also fed of Chironomidae at a depth > 10 m (Figure 4). This feeding behaviour of *O. agassizii* was also described for saline ecosystems populations of the southern of Altiplano (Chile), where a wide diet was found (Guzmán & Sielfeld, 2009). Nevertheless, even though such behaviour was also reported for this species in Lake Titicaca (Lauzanne, 1992), *O. agassizii* showed to taking advantage of prey abundance in the habitat.

According to Monroy et al., (2014), *O. agassizii* is an omnivorous species, and like other Cyprinodontiform species (Kalogianni *et al.*, 2010; Alcaraz *et al.*, 2015) it is not possible to generalize the feeding pattern for this species throughout the lake without considering other influencing factors (Saikia, 2016; Yoğurtçuoğlu *et al.*, 2018). A clear influence of seasonality and depth was observed in the results, which has influenced by feeding habits changing the proportions and importance of the prey (Table 4). Amphipods, zooplankton and ostracods seem to be the main food resource for *O. agassizii* across the region (Guzmán & Sielfeld, 2009; Maldonado *et al.*, 2009). For this reason, it could be mentioned that *O. agassizii* is a species capable of adapting its diet to the existing resources in the habitat, modifying it according to the season and availability.

*O. mulleri*, also classified as omnivorous (Monroy *et al.*, 2014) is described, by only observations as a species that bases its diet on molluscs and ostracods (Lauzanne, 1992; Vila *et al.*, 2007). However, like the other *Orestias* studied, *O. mulleri* fed mainly on amphipods, with a frequency higher than 50% for both seasons. Molluscs (Hidrobiidae) were the secondary prey during rainy season, and ostracods (Cyprididae) and Chironomidae for dry season (Figure 4 and 5). There is a lack of knowledge on the feeding habits of this species, so this work is the first to contain detailed information on their diet.

Although *Orestias* diet does not showed changes in its composition between depths and seasons, it certainly does the prey abundances in the gut content. It is possible to mention that *Orestias* have an opportunistic diet, consuming the prey with higher abundance in the ecosystem. Low variation in prey richness, the reduced use of trophic resource and the observed overlap showed a certain preference for amphipods in *Orestias* diet. Therefore, the usage increase or decrease of one resource may be due to spatial segregation, something suggested for these native species of the lake (Monroy *et al.*, 2014). During this study, the *Orestias* fish showed that all of them can feed on the same prey, even overlapping their diets (Figure 6). Diet overlapping increased during dry season (Figure 7), where the water temperature decreased a few degrees (Dejoux & Iltis, 1992). Temperature decreasing could influence the prey availability, or could reduce the effort for searching food by these fish.

Amphipods are the most exploited food resource, not only for native fish populations in Lake Titicaca, but also for introduced fish (*Odontesthes bonariensis* and *Oncorhynchus mykiss*) (Vaux *et al.*, 1988; Vila *et al.*, 2007). It is worth mentioning that during this investigation three possible species of amphipods were identified: *Hyalella* cf. *cuprea, Hyalella* cf. *latimanus*, and in deep zones (> 10 m) *Hyalella* cf. *longipes. Hyalella* cf. *cuprea* and *Hyalella* cf. *longipes* were observed in habitat and in the gut contents of *O. mulleri*. The most abundant in the habitat as well in the gut contents was *Hyalella* cf. *cuprea*.

Amphipods and molluscs represent an important component in the diet of other vertebrates in Lake Titicaca. For example, Titicaca’s water frog (*Telmatobius culeus*) (Muñoz-Saravia, 2018) diet is based on amphipods an molluscs, similar to *O. luteus* and *O. mulleri*. Nutritional values of amphipods show an important energy contribution (crude protein = 43%, gross energy = 13 kJ / g) (Muñoz-Saravia, 2018). This, added to the great abundance of this group in its habitat, could be the reason why most of the aquatic vertebrates of this ecosystem feed on this resource. However, it is not the same with molluscs (Hydrobiidae), whose nutritional contribution is clearly lower than other groups (crude protein = 15.6%, gross energy = 3.4 kJ / g) (For more information on nutritional composition in the diet of Titicaca water frog refer to Muñoz-Saravia, 2018). Although molluscs were an important component of *Orestias* diet, these do not seem to have an important nutritional contribution because molluscs do not undergo any change as digestion progress, making their identification even easier in the gut content (Hyslop, 1980; Baker *et al.*, 2014).

Some molluscs even survive passing through the digestive tract of the fish. Lazzaro (1987) reports that the ostracods *Cypriodopsis vidua* (Family Cyprididae) survives passing through the intestine of the sunfish (*Lepomis macrochirus*, Rafinesque, 1810) leading a negative selectivity to this prey (Vinyard & O’brien, 1976).

Planktivorous fish developed a strategy to avoid the deficiency in feeding due to the low digestibility caused by molluscs increasing the intake of these preys (Lazzaro, 1987). Muñoz-Saravia (2018) suggests that, in the case of Titicaca’s water frog, feeding of molluscs can help to shred the amphipods exoskeleton. Another possibility is that feeding of molluscs may delay the passage of food through the intestine giving a higher digestion time for the nutrient assimilation. This strategy could be the same for *Orestias* due to the high abundance of molluscs in its diet, especially for fish associated to the bottoms of the lake (*O. luteus* and *O. mulleri*). Importance of amphipods and molluscs in *Orestias* diet highlights the need for more studies focused on the nutritional profile of these fish in wild conditions, as well as the nutritional contribution they provide.

In conclusion, *Orestias* species inhabiting the Toke Pucuro bay of Lake Titicaca base their diet on amphipods and molluscs. The observed depth-related changes support spatial segregation among these fish, nevertheless, the change in prey abundance of *Orestias* diet is more influenced by seasonality. Based on diet composition *O. luteus* and *O. mulleri* are invertivore species. *O. mulleri* has a greater diet breadth in relation to the prey richness on the habitat. On the other hand, *O. agassizii* showed to be an opportunistic omnivorous that feeds of the resource that has the highest abundance, shifting its feeding habits as the depth in the habitat increases. Due the use of the same trophic resource, these three *Orestias* species compete against themselves, which is more evident during the dry season. It is necessary to guide future researches on these species to analyse their food and nutritional requirements, considering the possibility of breeding them in captivity, which could reduce the exploitation of wild populations already affected by local overfishing.

## ACKNOWLEDGMENTS

Funding for this study came from the *“Erika Geyger” grants for young researchers* from Instituto de Ecología-UMSA and FUNDECO (Fundación para el Desarrollo de la Ecología). I am grateful to Rogelio Alcón (fisherman), Ariel Kallisaya, Oscar Ayala, Jorge Molina-Rodríguez, Carla Ibáñez, Oscar Rollano, Valeria Palma and Arturo Muñoz-Saravia for their assistance, help with the literature review and discussion on the relevance of feeding ecology of native species.

